# Cell Cycle-Dependent TICRR/TRESLIN and MTBP Chromatin Binding Mechanisms and Patterns

**DOI:** 10.1101/2024.02.02.578516

**Authors:** Tyler D. Noble, Courtney G. Sansam, Kimberlie A. Wittig, Blanka Majchrzycka, Christopher L. Sansam

## Abstract

The selection of replication origins is a defining characteristic of DNA replication in eukaryotes, yet its mechanism in humans has not been well-defined. In this study, we use Cut&Run to examine genomic binding locations for TICRR/TRESLIN and MTBP, the human orthologs for the yeast DNA replication initiation factors Sld3 and Sld7. We mapped TRESLIN and MTBP binding in HCT116 colorectal cancer cells using asynchronous and G1 synchronized populations. Our data show that TRESLIN and MTBP binding patterns are more defined in a G1 synchronized population compared to asynchronously cycling cells. We also examined whether TRESLIN and MTBP are dependent on one another for binding. Our data suggest MTBP is dependent on TRESLIN for proper association with chromatin during G1 but not S phase. Finally, we asked whether TRESLIN and MTBP binding to chromatin requires licensed origins. Using cell lines with a non-degradable inducible Geminin to inhibit licensing, we show TRESLIN and MTBP binding does not require loaded MCMs. Altogether, our Cut&Run data provides evidence for a chromatin binding mechanism of TRESLIN-MTBP during G1 that is dependent on TRESLIN and does not require interactions with licensed origins.

## Introduction

DNA replication is a fundamental process in cell biology, essential for faithfully transmitting genetic information from one generation to the next. The initiation of DNA replication is a highly regulated and precisely orchestrated process that ensures the accurate duplication of the genome before cell division. At the heart of this process are intricate protein-protein and protein-DNA interactions that coordinate the assembly and activation of the replication machinery. The initiation of DNA replication is regulated at three stages: origin licensing, selection, and firing. Licensing is the assembly of pre-replicative complexes (pre-RC) onto genomic sites where DNA replication may later initiate. Although the number of licensed origins may set the upper limit of replication initiation, it does not generally determine DNA replication rates because an excess number of pre-RCs are loaded onto potential origins than will ever be used (1, 2, 3, 4). Instead, the replication initiation rate is controlled at the origin firing step and is limited by the availability of origin firing factors (4, 5, 6).

Our current understanding of these processes has been primarily derived from studies in yeast. In *S. cerevisiae* the timing of origin firing is established during G1 after origin licensing. This selection of early firing origins involves the recruitment of the homologs Sld3, Sld7, and CDC45 to DDK-phosphorylated MCM2-7 at origins (7, 8). The rate-limiting step in this process is the recruitment of DDK by forkhead transcription factors or a kinetochore protein to early origin sites (6, 9). The binding of Sld3-Sld7 serves as a critical mark, designating these locations as early firing origins, ensuring that replication begins precisely when and where needed (7).

While these findings in yeast have provided valuable insights into the fundamental principles of DNA replication initiation, how this process unfolds in human cells is less clear. TRESLIN (also known as TICRR) and MTBP are the functional orthologs of the yeast Sld3/Sld7. Despite functional overlaps between TRESLIN/MTBP and Sld3/Sld7, these factors have evolved to accommodate distinct mechanisms of replication control (10, 11, 12, 13, 14). TRESLIN and MTBP are larger and bear little sequence similarity to Sld3 and Sld7 (15, 16, 17, 18). Until recently, the genomic binding of human-origin firing factors was unmapped. Kumagai et al. published that MTBP binds in human DLD1 cells to early-initiating sites, suggesting that recruitment of MTBP to those sites affects origin firing timing and efficiency (19). They also show that truncation of MTBP to remove a C-terminal G-quadruplex DNA binding sequence reduces its binding to the genome (19).

Despite these advances, key questions remain unanswered. We still lack a clear picture of where, when, and how TRESLIN and MTBP bind to the genome. Binding maps of TRESLIN have not been reported. It is unknown whether TRESLIN and MTBP bind to the same sites and whether their binding to these sites is interdependent. The published map of MTBP binding was derived from asynchronously cycling DLD1 cells, so it is unknown whether it binds to the same sites in other cell lines or at specific cell cycle phases. Finally, although work from Kumagai et al. suggests a model where MTBP is recruited to origins independent of the loading of the MCM2-7 complex, the role of origin licensing in the recruitment of TRESLIN or MTBP to origins in human cells has not been determined.

This study seeks to explore the roles of MTBP and TRESLIN in human cells, shedding light on their binding locations, interactions, and potential differences in mechanisms. We employed Cut&Run to map TRESLIN and MTBP binding locations genome-wide. Here we show that binding of TRESLIN and MTBP is cell cycle stage-specific and this binding is independent of origin licensing. These results suggest a model that diverges from that seen in budding yeast. Additionally, we present evidence for a role for TRESLIN in the recruitment of the TRESLIN/MTBP complex to chromatin during G1.

## Results

### Reproducible binding of TRESLIN and MTBP to origins and early replicating domains

We aimed to define genomic binding locations for TRESLIN and its partner MTBP and to determine whether they bound to the same sites. Initial attempts using antibodies against endogenous proteins failed due to high background signal. Using CRISPR-Cas9 we tagged all alleles of TRESLIN or MTBP in HCT116 cells with monomeric (m)Clover which allowed use of an antibody against GFP for Cut&Run (20). We previously showed that our mClover tagged cell lines proliferate normally indicating that the tag does not impair normal function of the proteins (20, 21). In total, we generated Cut&Run data from 46 samples for this study, and we used all samples to identify consistent binding regions. A peak was considered reproducible if detected in at least six Cut&Run experiments for each protein. Our analysis revealed shared binding sites between TRESLIN and MTBP, and we detected few sites occupied by either protein alone (Fig. 1A, C, D).

**Fig 1.**
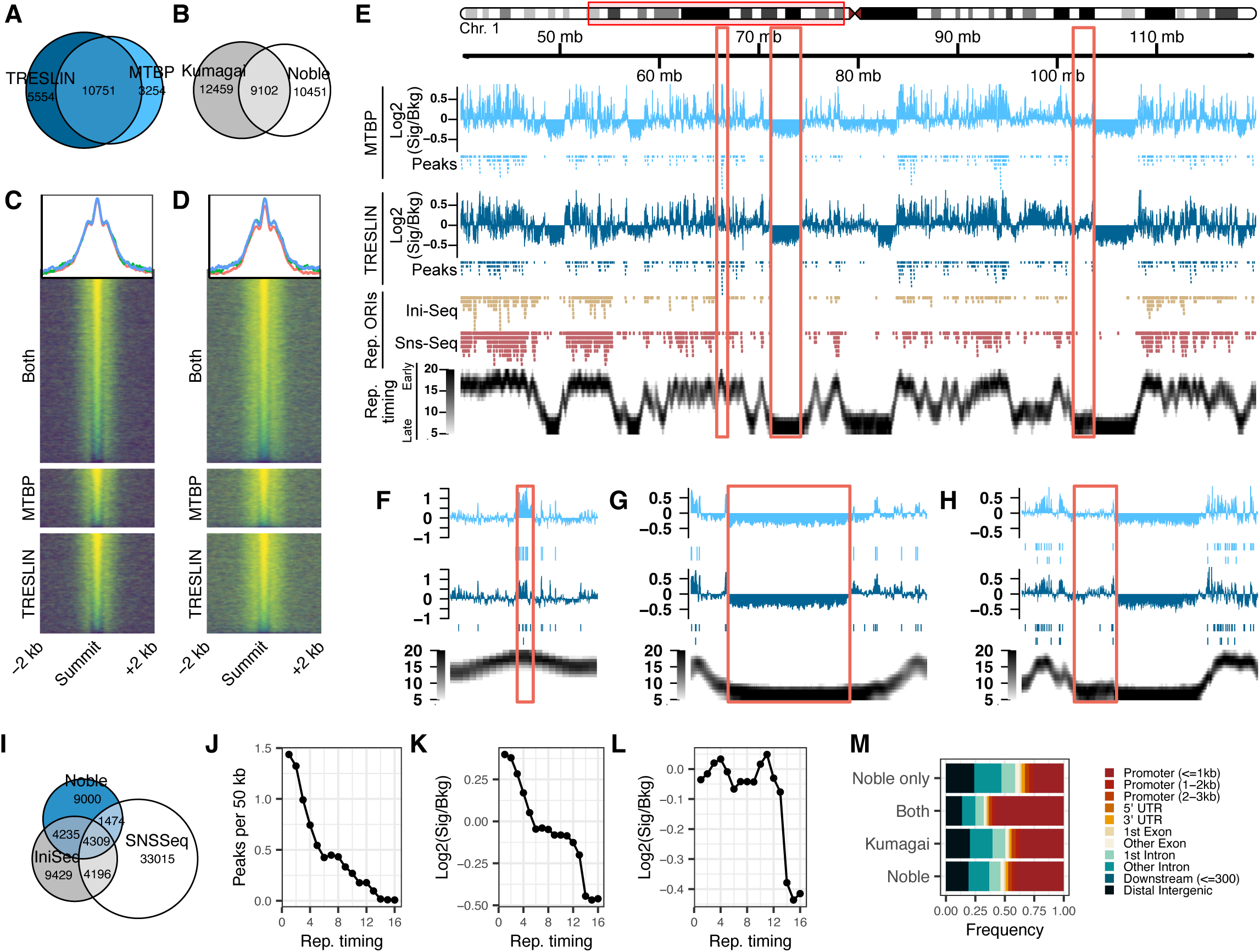
TRESLIN and MTBP co-occupy replication origins and early-replicating regions in HCT116 cells. (A). Cut&Run experiments were conducted on HCT116 cells expressing MTBP or TRESLIN genes tagged with mClover. Peaks were called with MACS2 and were designated highly reproducible if found in at least 6 Cut&Run experiments. Euler plot illustrates the extensive overlap of highly reproducible TRESLIN and MTBP peaks. (B) Euler plot shows that MTBP/TRESLIN peaks in HCT116 cells significant overlap peaks from published MTBP Cut&Run data, in which MTBP in DLD1 cells was endogenously FLAG-tagged(19). (C) Heat plots display MTBP Cut&Run signal (Log2 fold-change over background) across high-confidence peaks. (D) As in C, heat plots show TRESLIN signal across high-confidence peaks. (E) Genome track set for human chromosome 1 shows that the istribution of MTBP and TRESLIN binding sites is associated with early replicating domains and DNA replication origins. The following are displayed: Log2Fold Change signal for MTBP (light blue) and TRESLIN (dark blue), along with locations of peaks, DNA replication origins (Rep. ORIs), and high-resolution Repli-Seq data(24, 50, 51). (F) Left highlighted region in E shows a replication timing peak with abundant TRESLIN/MTBP signal. (G) Central region in E depicts late replicating areas with decreased TRESLIN/MTBP signal. (H) Right region in E. A portion of the late-replicating region that replicates earlier and has increase TRESLIN/MTBP binding is highlighted. (I) Euler plot illustrates the overlap of highly reproducible TRESLIN/MTBP peaks with Ini-Seq and Sns-Seq origins(50, 51). (J) Counts of TRESLIN/MTBP peaks in 50 kb genome segments across 16 replication timing fractions, ranging from early to late replication. (K) Mean Cut&Run Log2FC signal over background for MTBP (K) or TRESLIN (L) plotted for each replication timing fraction reveals that MTBP and TRESLIN binding correlates with replication timing genome-wide. (M) Analysis of TRESLIN/MTBP peak locations relative to gene features reveals frequent presence in gene promoters, especially in peaks also found in DLD1 cells(19).

Kumagai et al. demonstrated MTBP bound to specific regions in human DLD1 cells by tagging MTBP with FLAG and employing Cut&Run (22). When we compared our TRESLIN/MTBP Cut&Run peak data with the previously published MTBP DLD-1 Cut&Run data, we observed a significant overlap in binding sites between the two datasets (Fig. 1B). Our analysis of genomic features associated with TRESLIN/MTBP peaks revealed that both proteins frequently bound to gene promoters in HCT116 cells, resembling the binding patterns of MTBP in DLD1 cells as reported by Kumagai et al. (22). Peaks identified in both studies exhibited the strongest association with gene promoters, while peaks exclusive to HCT116 cells were in other gene or intergenic regions (Fig. 1M). The consistency between the MTBP map in DLD1 cells and the TRESLIN/MTBP map in HCT116 cells underscores the robustness of both data sets and highlights commonalities in the binding patterns of these proteins in different cell contexts.

We next asked if our TRESLIN and MTBP binding would be enriched in early replicating domains and at replication origins, as this has been reported for MTBP in DLD-1 cells (19). To this end, we compared regions of TRESLIN and MTBP binding to Ini-Seq, SNS-Seq, and high resolution Repli-Seq datasets (Fig. 1E-L) (23, 24, 25). Early-replicating regions have the highest density of TRESLIN/MTBP peaks, and late-replicating domains have very few (Fig. 1E, J). TRESLIN and MTBP also appear to be diffusely bound in early replicating regions outside of peaks, while there is an obvious depletion in diffuse TRESLIN/MTBP signal from late-replicating domains (Fig. 1E-H). Interestingly, late-replicating domains are often split into late and mid/late replicating segments, and there is a clear shift in the level of diffuse TRESLIN/MTBP signal at that transition (Fig. 1E, G-H). At sites where there is a sharp early replicating peak, which likely represents an efficient initiation zone, there are typically one or more TRESLIN/MTBP peaks and broad enrichment of signal from both proteins (Fig. 1E, F). Although narrow TRESLIN/MTBP peaks are enriched mainly in early replicating genomic regions, diffuse binding of both proteins extends into regions of the genome replicated during the middle of S phase (Fig. K, L). This correlation between replication timing and TRESLIN-MTBP binding suggests a functional role for origin selection. Sites of binding that were common between TRESLIN and MTBP also showed substantial overlap with Ini-Seq peaks from EJ30 cells from and SNS-Seq peaks from HCT116 cells (Fig. 1I) (25, 26).

### TRESLIN and MTBP bind to potential replication origins during G1

Given that Sld3/Sld7 binds to origins before S phase begins, we next sought to determine if TRESLIN and MTBP binding was enriched at specific sites in G1 compared to an asynchronously cycling population. For this, we synchronized cells using a nocodazole arrest followed by mitotic shake-off and release to obtain cells enriched in G1 (Fig. 2A). To confirm enrichment of cells in G1 phase, we assessed the DNA content by flow cytometry prior to performing Cut&Run (Fig. 2B, and 2C). We completed three Cut&Run replicates of each for both TRESLIN and MTBP cells in G1 and cycling cells. We found that there was an increased signal in G1 for both TRESLIN and MTBP compared to cycling samples by comparing background-normalized Cut&Run signal across consensus peaks (Fig. 2D-G). The extent of the difference in signal in peaks between G1 and cycling cells suggests that like in yeast, TRESLIN and MTBP bind to origins during G1. However, the reduced Cut&Run signal in cycling cells compared to G1, indicates that both proteins are released from the G1 binding sites as they progress into S phase.

**Fig 2.**
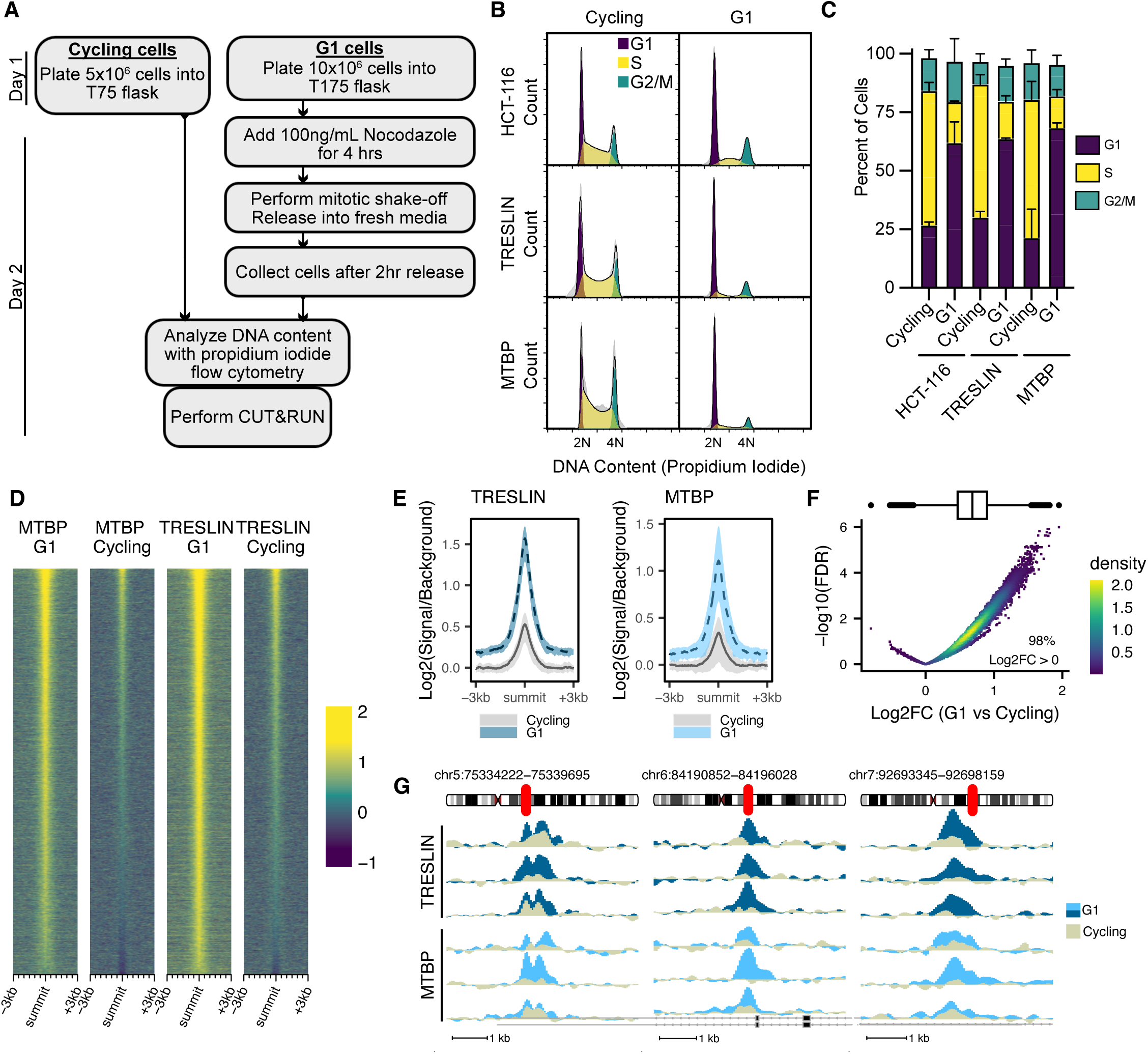
TRESLIN and MTBP enriched in genomic peaks during the G1 phase of the cell cycle. (A) Schematic of the experimental design comparing TRESLIN and MTBP localization in freely cycling cells and G1-phase cells. Freely cycling cells underwent natural cell cycle progression, while G1 cells were collected post-Nocodazole-induced mitotic arrest. (B) DNA content histograms from a single replicate, illustrating cell cycle distribution in cycling and G1-enriched cells. Histograms highlight cell cycle phases (color-shaded) as estimated by the Dean-Jett-Fox algorithm, with parental HCT-116 cells serving as background controls. (C) Flow cytometry-based quantification of cell cycle phase percentages in two independent replicates. (D) Heat plot of background-normalized Cut&Run signals (Log2FC) across MACS2-identified peaks in at least two samples. (E) Aggregate plot of Log2-fold changes in background-normalized MTBP or TRESLIN signals in consensus peaks relative to HCT-116 cells across three replicates. Lines represent mean values, with ribbons indicating the range for these replicates. (F) Volcano plot of DESeq2 analysis showing FDR versus changes in MTBP and TRESLIN signals at consensus peaks, comparing G1 to cycling cells. Signals were predominantly higher in G1 cells. (G) Background-normalized signals across three representative ranges, indicating reduced peaks in cycling compared to G1 cells. Data for TRESLIN and MTBP are shown separately for each of the three replicates.

### A role for TRESLIN in the localization of TRESLIN-MTBP complex to chromatin during G1

TRESLIN and MTBP are known binding partners but the extent of their interdependency for proper DNA binding is unknown (10, 12, 27). We used flow cytometry to assess both total protein and chromatin-bound protein for TRESLIN and MTBP under conditions of siRNA knockdown. Total protein levels for MTBP did not significantly change when cells were treated with siTRESLIN (Fig. 3A-B). Conversely, total levels of TRESLIN showed a significant decrease during G1 when cells were transfected with siMTBP (Fig. 3A-B).

**Fig 3.**
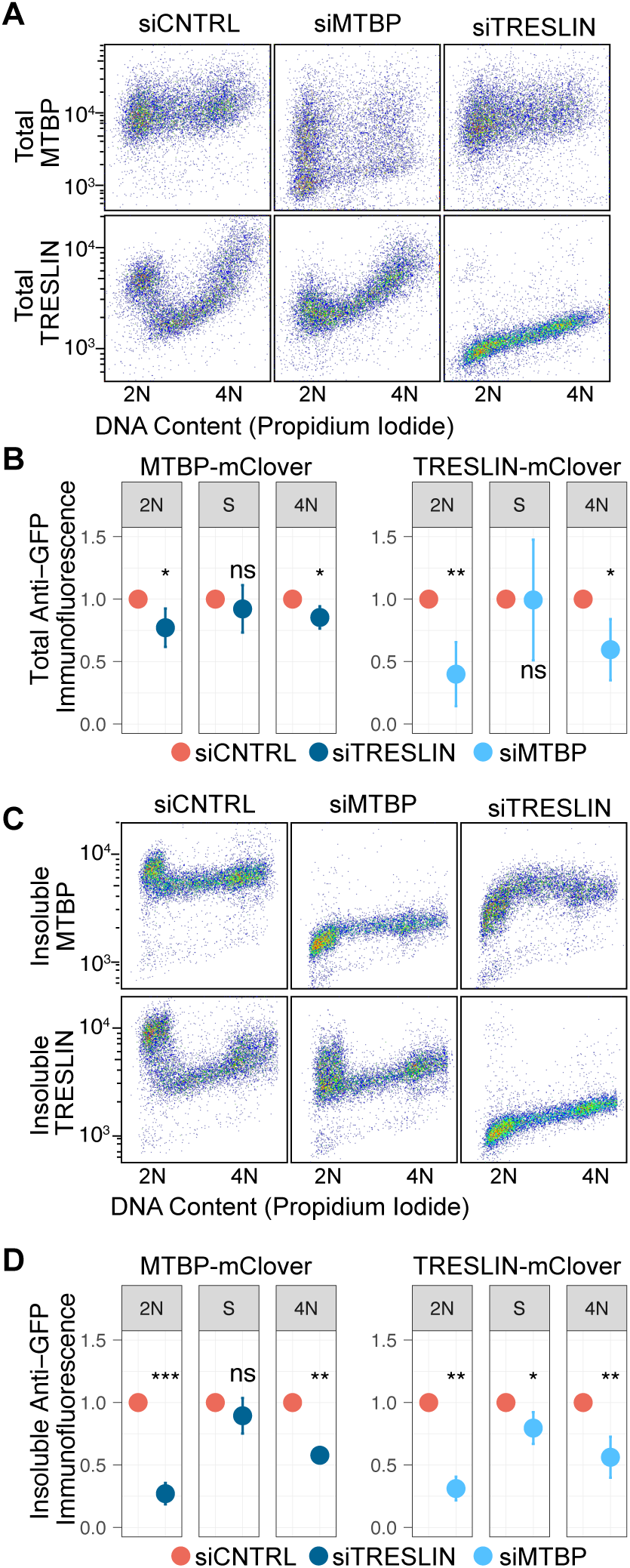
Binding of MTBP to chromatin requires TRESLIN in G1 but not in S. (A) Protein levels of MTBP-mClover and TRESLIN-mClover in HCT116 cells were measured via flow cytometry using an anti-GFP antibody, alongside DNA content. This analysis was conducted 24 hours after siRNA transfection, representing a single experimental replicate. (B) Quantification of the data in panel A involved background-subtracting individual cell anti-GFP immunofluorescence signals, averaging signals for cells in 2N, early S, or 4N DNA content gates, and normalizing values from siTRESLIN or siMTBP transfected cells to siCNTRL. The means of these normalized values across three replicates are shown with error bars indicating 95% confidence intervals. Asterisks indicate significance levels from t-tests comparing each sample to its respective siCNTRL. (C) and (D) replicate (A) and (B), respectively, with the exception that soluble protein was pre-extracted before fixation. Consequently, immunofluorescence signals represent insoluble TRESLIN-mClover or MTBP-mClover protein.

Next, we looked at chromatin-bound or insoluble levels of both proteins. Not surprisingly, insoluble levels of TRESLIN closely mirrored its total levels; both exhibited reductions in response to siRNA targeting MTBP (Fig. 3C-D). Strikingly, siRNA knockdown of TRESLIN caused a strong reduction in chromatin-bound MTBP during G1 and a milder reduction during G2 (Fig. 3C-D), even though total levels of MTBP were not substantially reduced. These results show that during G1, MTBP is dependent on the presence of TRESLIN to bind chromatin, suggesting a TRESLIN-dependent mechanism for the localization of TRESLIN-MTBP to potential origins. Interestingly, the level of chromatin-bound MTBP during S phase was not affected by TRESLIN knockdown.

### Licensing is not a requirement for TRESLIN and MTBP binding to chromatin

Since yeast Sld3 and Sld7 are known to bind origins via MCM we next asked whether TRESLIN and MTBP similarly require MCMs for the genomic binding mapped with Cut&Run (8, 28). We constructed cell lines that prevent loading of MCMs using a doxycycline(dox)-inducible G1-stabilized GEMININ (T25D) (Supplemental Fig. 1) (29). Expression of the GEMININ(T25D) transgene causes an accumulation of cells in G1 (Supplemental Fig. 1). Therefore, all cells were synchronized in G1 phase by initially treating them with thymidine, followed by mimosine treatment while concurrently administering dox to prevent licensing (Fig. 4A). The decrease in MCMs loading was verified by flow cytometry using antibody against MCM7 (Fig. 4B). TRESLIN and MTBP binding was assessed in two ways: immuno-flow cytometry measurement of insoluble protein and binding mapped using Cut&Run with fixation. Immuno-flow cytometry showed binding for both TRESLIN and MTBP persisted after inhibition of licensing (Fig. 4B and Supplementary Fig. 1). Licensing inhibition did not affect TRESLIN or MTBP enrichment in peaks. Heat plots of spike-in normalized Cut&Run signals for TRESLIN or MTBP across consensus MACS2 peaks revealed consistent signal levels in all samples, whether they expressed Geminin(T25D) or not (Fig. 4C, D, F). The inhibition of licensing also had no impact on the overall enrichment in early replicating regions compared to late replicating regions, as the difference in the overall Cut&Run signal between early and late replicating genomic windows remained unaltered by the expression of Geminin(T25D) (Fig. 4E). Interestingly, MTBP binding exhibited a slight reduction in the latest timing domains in the presence of Geminin(T25D). Subsequent experiments will be required to discern whether this alteration in MTBP binding is of biological significance or if it stems from variability within the more challenging-to-map regions of compact heterochromatin (Fig. 4E). Taken together these data demonstrate binding of TRESLIN and MTBP in sharp peaks or broadly across early- and mid-replicating domains is not dependent on origin licensing, suggesting a model of binding for initiation factors that diverges from yeast models.

**Fig 4.**
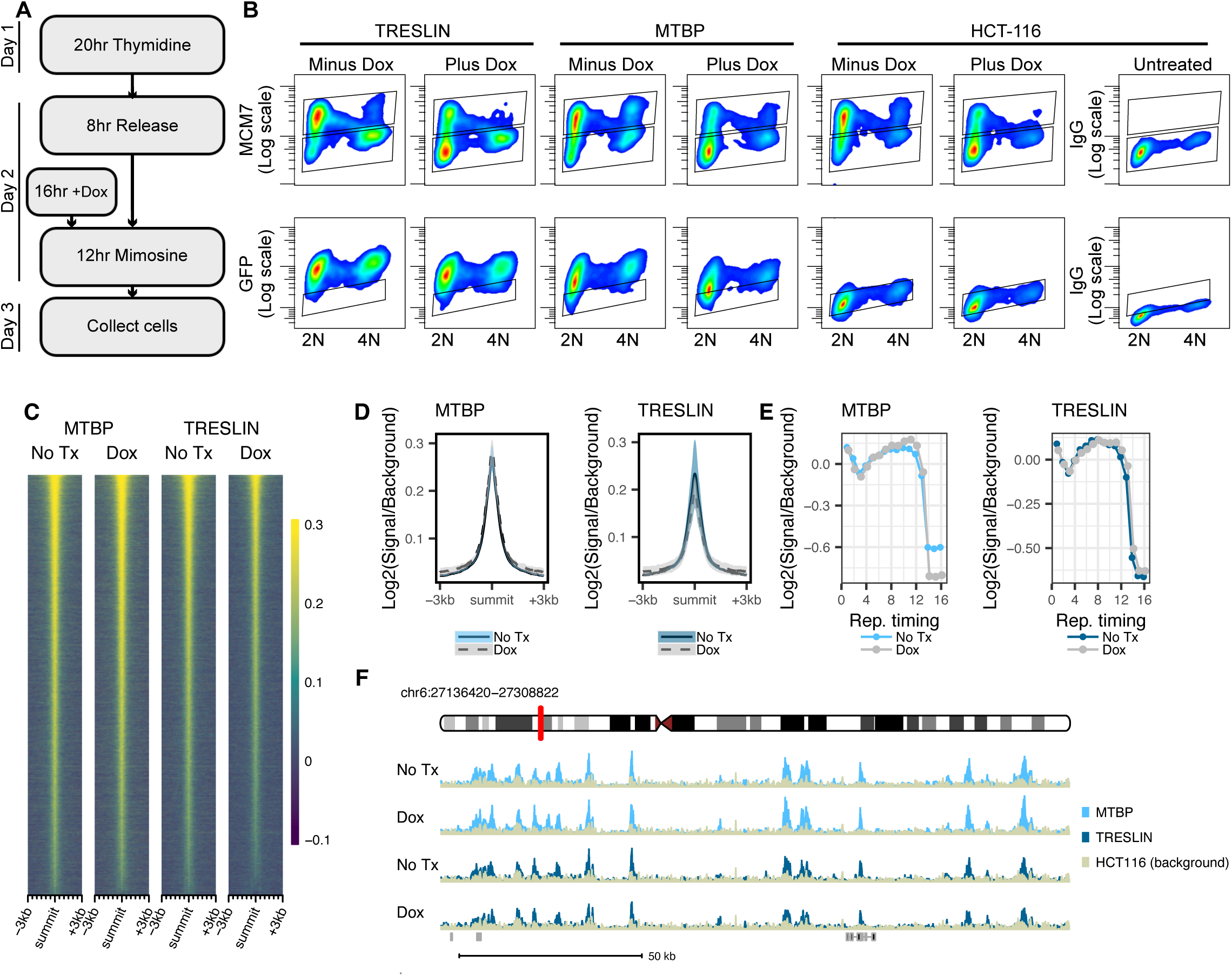
Licensing is not required for TRESLIN and MTBP binding to genomic loci and early replicating regions. A) Diagram of cell synchronization strategy and induction of Geminin transgene expression to inhibit loading of MCMs. B) Insoluble flow cytometry showing level of chromatin bound MCM7 and TRESLIN-mClover or MTBP-mClover in tagged (TRESLIN or MTBP) or untagged (HCT-116) cells. Anti-MCM7, anti-GFP, or anti-IgG Immunofluorescence signal vs DNA content is plotted to show the level of each protein across the cell cycle. Treatment with doxycycline for 16 hours was sufficient to reduce levels of chromatin bound MCM7 in 2N cells, but it did not affect TRESLIN or MTBP binding. C) Heat plots showing Cut&Run signal over peaks called with MACS2 with or without doxycycline treatment over reproducible TRESLIN or MTBP peaks. D) Aggregate plots of Log2fold signal over HCT116 parental untagged background. Plots show two replicates for each. E) Log2fold signal of Cut&Run for TRESLIN and MTBP plotted against replication timing from high resolution Repli-Seq data(24) shows that licensing inhibition does not affect TRESLIN/MTBP enrichment in early replicating regions F) Example genomic range of peaks present in samples treated with or without doxycycline. Scale = 50kb.

## Discussion

Binding of initiation factors is a key step in origin selection, but in human cells the specific mechanisms that determine which origins are selected remain unclear. Here we sought to define binding locations and conditions of binding for TRESLIN and MTBP to elucidate their role in the selection and firing of origins.

TRESLIN and MTBP binding was found to be localized together at the same genomic sites. When we compared our asynchronous MTBP loci to those published in DLD-1 cells we found overlap between the two data sets in active regions of the genome i.e. upstream of promoters, while differences in binding were largely in intergenic or intronic regions. As HCT116 and DLD-1 are both colorectal cancer cell lines it was expected that MTBP loci may be similar. Future examination of cell lines from different lineages will be helpful in establishing the direct role of these initiation factors and how they relate to the efficient origin sites that have been mapped by sequencing and optical approaches (23, 30, 31).

According to the yeast model of Sld3 and Sld7, the binding of these initiation factors occurs during G1 to prepare for the upcoming S phase (7). In our study with HCT116 cells, we observed that TRESLIN and MTBP exhibited the highest enrichment in their chromatin binding during the G1 phase, as compared to an asynchronous population. This finding suggests that these human proteins may share a similar role in promoting firing at specific origins by binding during G1. Our results also revealed that the presence of loaded MCM complexes was not a prerequisite for TRESLIN and MTBP to bind chromatin during G1. This implies that TRESLIN/MTBP are recruited to early origins through a mechanism distinct from binding to phosphorylated MCMs. Intriguingly, prior research has shown that MCM-independent chromatin association of TRESLIN also occurs in X. laevis extracts, though the precise underlying mechanism remains unidentified (12).To further elucidate the orchestration and regulation of this binding, follow-up experiments will be essential to determine the requirements of DDK and CDK activity for the observed localization patterns observed via Cut&Run.

Since it is known that TRESLIN and MTBP are binding partners (10, 27) we hypothesized that their binding would be interdependent. Interestingly, we found that MTBP binding was affected by siRNA knockdown of TRESLIN. It has previously been shown that MTBP has the ability to bind chromatin directly through its C-terminus (10, 19), but no direct binding function has been found for TRESLIN. We consider the possibility of undefined binding partners for the TRESLIN-MTBP complex that may explain our findings through additional proteins interacting with TRESLIN. Another possible explanation is the TRESLIN-BRD4 interaction and potential recruitment to chromatin via acetylated histones (32). We propose a model where TRESLIN/MTBP are recruited to chromatin during G1 in a TRESLIN-dependent manner. Intriguingly, MTBP binding during S phase was largely unaffected to TRESLIN knockdown. This is consistent with our previous work showing little change in chromatin bound MTBP levels when TRESLIN levels drop at the G1/S transition, and it suggests that MTBP may bind to potential origins during S phase without TRESLIN(20).

Taken together, the data presented here illustrate conditions of binding for TRESLIN and MTBP. By mapping the binding of these two interaction partners separately, we have provided unequivocal evidence for their co-localization within the genome. Additionally, we have revealed potential differences in the mechanisms of origin selection in humans compared to yeast that offer exciting avenues to explore. Ultimately a strategy that employs perturbation of TRESLIN or MTBP binding followed by direct measurements of origin firing through sequencing or optical replication mapping approaches will be needed to fully define the mechanisms considered here (23, 30, 31). The data from this study provides new insights into the binding of TRESLIN and MTBP that will be useful for defining the key steps of origin selection in human cells.

## Materials and Methods

### Cut&Run

Cut&Run was performed as described by Skene et al., 2018 using an anti-GFP antibody from Rockland Immunochemical (cat 600-401-215S). For standard Cut&Run, each sample used 100,000 cells and protein A-MNase digest step used 40min incubation. For fixation, Cut&Run each sample used 300,000 cells and included a modified wash buffer and fixation steps adapted from CUTANA Cross-linking Protocol v1.9 from Epicypher. Samples were fixed with either 0.1 or 1% fresh paraformaldehyde for 2 min and quenched with 2.5M Glycine at 1:20 dilution before proceeding to wash steps. Overnight incubation at 55C with 2uL proteinase-K (20mg/mL) and 1.6uL 10% SDS per sample was used to reverse cross-links before DNA purification. For asynchronous Cut&Run experiments HCT-116 cells were plated at 2.5×10^^6^ in 10 cm^2^ dishes and collected after 48hrs.

### Flow Cytometry

Chromatin-binding immuno-flow cytometry experiments were performed as described in Wittig et al., 2021. For MCM7 and mClover evaluation, MCM7 antibody was purchased from Santa-Cruz (Cat# sc-56324) and GFP antibody was purchased from Rockland Immunochemicals (cat 600-401-215S). DNA content assessed by flow cytometry was performed by resuspending cell pellets in a DNA staining buffer (0.1mg/mL propidium iodide, 0.1% Na Citrate, 0.02mg/mL RNase A, 0.3% NP-40, ddH2O) and incubating at room temperature for at least 30min before analysis.

### Cell Culture

HCT-116 cells were purchased from ATCC and cultured in McCoy’s 5A medium (Corning 10-050-CV) with 10% Fetal Bovine Serum.

### Cell Cycle Synchronization

For G1 synchronized experiments, cells were plated at a density of 10×10^^6^ cells into T175 flask. The next day, 100ng/mL of Nocodazole (Sigma M1404) was added to the media for 4 hours followed by a mitotic shake-off. Cells were washed 4X in 50mL 1xPBS and replated into fresh McCoy’s 5A medium containing 10% FBS for 2 hours. For an accompanying asynchronous sample, 5×10^^6^ HCT-116 cells were plated into a T75 flask and collected in parallel. DNA content was analyzed with propidium iodide using flow cytometry.

### siRNA treatment

Cells were plated at 250,000 cells per well in a 6-well dish and allowed to recover overnight. siRNA transfection was carried out using RNAiMAX reagent (ThermoFisher Scientific 13778075). siRNA MTBP (Dharmacon, M-013953-01-0010), siRNA TRESLIN (Life Technologies, (<u>Kumagai et al. 2010</u>), and siControl (Life Technologies, 12935110) were used at 20nM final concentration and incubated for 24 hours before collecting cells for flow cytometry.

### Geminin Cell Line Construction

Doxycycline-inducible Geminin cell lines were constructed in our TICRR-mClover, MTBP-mClover, and HCT-116 cell lines. Geminin stabilization is regulated by a phosphorylation on Threonine 25, so T25 was mutated to Aspartate (T25D) to prevent its degradation (33). To construct the GemininT25D Piggybac transposon plasmid, a synthetic DNA fragment was inserted into KpnI and SpeI-digested XLone-GFP using Gibson Cloning (34).

XLone-GFP was a gift from Xiaojun Lian (Addgene plasmid # 96930; http://n2t.net/addgene:96930; RRID:Addgene_96930) (35). pCMV-hyPBase was a gift from Allan Bradley via the Wellcome Trust Sanger Institute (36). The T25D-Geminin stable cell line was generated by transfecting (Mirus Bio) with two plasmids: XLone-GemininT25D and pCMV-hyPBase, a highly active PiggyBac transposase, using a 1:9 ratio (XLone:hyPBase) into the HCT-116 mClover_TICRR cell line. After 48 hours, the cells were split and selected in blasticidin (10ug/mL). Selected single cell clones were screened by DNA content flow cytometry for a G1 arrest phenotype after Doxycycline was added to the medium 24 hours to induce T25D Geminin expression.

### Licensing Inhibition Experiments

Cells were plated at a density of 700,000 per well in 6cm^2^ plates and allowed to recover overnight. Cells were then treated with 2mM Thymidine (Sigma T1895) for 22 hours followed by a wash in 1xPBS and a 6 hour release. At 2 hours into the release 1ug/mL doxycycline was added for a 16 hour dox treatment. After the 6 hours release 0.5mM mimosine was added for 12 hours. Cells were then collected and analyzed using chromatin-binding immune-flow cytometry to assess inhibition of MCM loading. Cut&Run was performed using fixation protocol and 300,000 cells per sample.

### Sequencing Library Preparation

Libraries for sequencing were prepared using KAPA Hyper Prep Kit (Ref# 7962347001-KK8502) following the manufacturer’s protocol. Adapters used were IDT for Illumina TruSeq UDI indexes (Ref# 20020178). Library amplification was performed using 13-16x cycles for Cut&Run samples and 3x cycles for Replication Timing samples. After post-amplification KAPA bead clean-up (Ref# 07983271001-KK8000) an additional 0.8x followed by 0.7x bead clean-up was performed as needed to remove adapter dimers. D1000 ScreenTape and Reagents (Agilent Ref# 5067-5582 and 5067-5583) for TapeStation were used to assess sample quality. KAPA library quantification kit (Ref# 0960298001-KK4854) was used to quantify samples on Roche LightCycler 480 II. Samples were sequenced on Illumina NovaSeq 6000 platform for PE150 sequencing.

### Cut&Run Analysis

The initial processing of Cut&Run sequencing data was conducted systematically using a tailored Snakemake pipeline(37, 38). First, we applied Trimmomatic to trim reads, excising bases with quality scores below 20 at both the leading and trailing ends, as well as internal bases with an average quality score lower than 15 within a 4-base pair window(39). To remove adapter remnants shorter than 6 bases, Cutadapt was employed with a quality cutoff of 20 and a mismatch rate of 20%(40). Reads shorter than 25 base pairs were filtered out after each trimming stage. Subsequently, we aligned the processed reads to the GRCh38 - hg38 human genome assembly using bowtie2(41). Default settings were utilized, with the inclusion of the --dovetail flag to permit dovetailed paired reads during alignment. Alignment processing and filtering was performed using bamtools, samtools, and sambamba to eliminate reads with a quality score less than 10, those with an unmapped mate pair, or duplicates(42, 43). Alignments or peaks overlapping problematic genomic regions (encode blacklist) were removed during downstream analysis(44). Genome coverage bigwig files were generated with deeptools bamCoverage(45). Peak calling was performed on individual replicates using macs2, applying an FDR threshold of 0.05 and a minimum fold change cutoff of 1(46). Master peaks were then called from alignments combined across all replicates for each set. To ensure equal representation of each replicate, alignments were randomly subsampled to match the read count of the replicate with the lowest coverage. Highly reproducible peaks were identified by calling master peaks from alignments across six asynchronously cycling cell replicates and assessing their overlap with peaks from other Cut&Run experiments. Peaks were deemed reproducible if they overlapped with those from at least 6 separate Cut&Run experiments. Custom R scripts, including the GenomicRanges package, were used for peak overlap analysis and downstream analyses(47). Differential binding analysis was conducted using DESeq2 within the Diffbind package(48, 49).

## Author Contributions

CLS and TDN wrote and prepared the manuscript. CLS, CGS, and TDN designed experiments. TDN and KAW generated flow cytometry data. TDN and CGS performed Cut&Run experiments. BM performed initial Cut&Run protocol validation. KAW constructed Geminin cell lines. CLS performed statistical analysis. CLS and TDN analyzed Cut&Run data.

## Acknowledgements

We thank the OMRF Clinical Genomics Center for sequencing services. We thank the Henikoff lab for providing an initial aliquot of pA-MNase. We thank Dr. Susannah Rankin and Dr. Allison Jevitt for purifying pAG-MNase. We thank Kevin Boyd and also OMRF’s Center for Biomedical Data Sciences for assistance with building the Cut&Run analysis pipeline.

## Funding

Funding provided by National Institutes of Health [R01GM121703] and Oklahoma Center for Adult Stem Cell Research. TDN received support from the John and Mildred Carson PhD Scholarship Fund awarded for the OMRF Pre-doctoral Scholarship.

Supplemental Fig 1: Timecourse of MCM depletion following doxycycline-induced Geminin expression. A) GemininT25D cell line with TRESLIN-mClover was treated with doxycycline for 4, 8, 12, 16, 20, and 24hr timepoints. Insoluble flow cytometry is shown for MCM7 and both total and insoluble flow cytometry are shown for TRESLIN. B) GemininT25D cell line with MTBP-mClover was treated with doxycycline for 4, 8, 12, 16, 20, and 24hr timepoints. Insoluble flow cytometry is shown for MCM7 and both total and insoluble flow cytometry are shown for MTBP. C) Quantification of insoluble flow cytometry showing median MCM7 signal compared to median TRESLIN signal from plots shown in A. D) Quantification of insoluble flow cytometry showing median MCM7 signal compared to median MTBP signal from plots shown in B.

## Notes

### Competing Interest Statement

The authors have declared no competing interest.

## References

1. Burkhart R, Schulte D, Hu D, Musahl C, Göhring F, Knippers R. Interactions of human nuclear proteins P1Mcm3 and P1Cdc46. Eur J Biochem. 1995;228(2):431–8.

2. Mahbubani HM, Chong JP, Chevalier S, Thömmes P, Blow JJ. Cell cycle regulation of the replication licensing system: involvement of a Cdk-dependent inhibitor. J Cell Biol. 1997;136(1):125–35.

3. Edwards MC, Tutter AV, Cvetic C, Gilbert CH, Prokhorova TA, Walter JC. MCM2-7 complexes bind chromatin in a distributed pattern surrounding the origin recognition complex in Xenopus egg extracts. J Biol Chem. 2002;277(36):33049–57.

4. Wong PG, Winter SL, Zaika E, Cao TV, Oguz U, Koomen JM, et al. Cdc45 limits replicon usage from a low density of preRCs in mammalian cells. PLoS One. 2011;6(3):e17533.

5. Collart C, Allen GE, Bradshaw CR, Smith JC, Zegerman P. Titration of four replication factors is essential for the Xenopus laevis midblastula transition. Science. 2013;341(6148):893-6.

6. Mantiero D, Mackenzie A, Donaldson A, Zegerman P. Limiting replication initiation factors execute the temporal programme of origin firing in budding yeast. Embo j. 2011;30(23):4805–14.

7. Tanaka S, Nakato R, Katou Y, Shirahige K, Araki H. Origin association of Sld3, Sld7, and Cdc45 proteins is a key step for determination of origin-firing timing. Curr Biol. 2011;21(24):2055–63.

8. Deegan TD, Yeeles JT, Diffley JF. Phosphopeptide binding by Sld3 links Dbf4-dependent kinase to MCM replicative helicase activation. Embo j. 2016;35(9):961–73.

9. Fang D, Lengronne A, Shi D, Forey R, Skrzypczak M, Ginalski K, et al. Dbf4 recruitment by forkhead transcription factors defines an upstream rate-limiting step in determining origin firing timing. Genes Dev. 2017;31(23-24):2405–15.

10. Kumagai A, Dunphy WG. MTBP, the partner of Treslin, contains a novel DNA-binding domain that is essential for proper initiation of DNA replication. Mol Biol Cell. 2017;28(22):2998–3012.

11. Sansam CG, Goins D, Siefert JC, Clowdus EA, Sansam CL. Cyclin-dependent kinase regulates the length of S phase through TICRR/TRESLIN phosphorylation. Genes Dev. 2015;29(5):555–66.

12. Volpi I, Gillespie PJ, Chadha GS, Blow JJ. The role of DDK and Treslin-MTBP in coordinating replication licensing and pre-initiation complex formation. Open Biol. 2021;11(10):210121.

13. Zaffar E, Ferreira P, Sanchez-Pulido L, Boos D. The Role of MTBP as a Replication Origin Firing Factor. Biology (Basel). 2022;11(6).

14. Zonderland G, Vanzo R, Gadi SA, Martín-Doncel E, Coscia F, Mund A, et al. The TRESLIN-MTBP complex couples completion of DNA replication with S/G2 transition. Molecular Cell. 2022;82(18):3350–65.e7.

15. Sanchez-Pulido L, Diffley JF, Ponting CP. Homology explains the functional similarities of Treslin/Ticrr and Sld3. Curr Biol. 2010;20(12):R509–10.

16. Kumagai A, Shevchenko A, Shevchenko A, Dunphy WG. Treslin collaborates with TopBP1 in triggering the initiation of DNA replication. Cell. 2010;140(3):349–59.

17. Sansam CL, Cruz NM, Danielian PS, Amsterdam A, Lau ML, Hopkins N, et al. A vertebrate gene, ticrr, is an essential checkpoint and replication regulator. Genes Dev. 2010;24(2):183–94.

18. Itou H, Muramatsu S, Shirakihara Y, Araki H. Crystal structure of the homology domain of the eukaryotic DNA replication proteins Sld3/Treslin. Structure. 2014;22(9):1341–7.

19. Kumagai A, Dunphy WG. Binding of the Treslin-MTBP Complex to Specific Regions of the Human Genome Promotes the Initiation of DNA Replication. Cell Rep. 2020;32(12):108178.

20. Wittig KA, Sansam CG, Noble TD, Goins D, Sansam CL. The CRL4DTL E3 ligase induces degradation of the DNA replication initiation factor TICRR/TRESLIN specifically during S phase. Nucleic Acids Res. 2021;49(18):10507–23.

21. Skene PJ, Henikoff S. An efficient targeted nuclease strategy for high-resolution mapping of DNA binding sites. eLife. 2017;6:e21856.

22. Kumagai A, Dunphy WG. Binding of the Treslin-MTBP Complex to Specific Regions of the Human Genome Promotes the Initiation of DNA Replication. Cell Rep. 2020;32(12):108178-.

23. Langley AR, Gräf S, Smith JC, Krude T. Genome-wide identification and characterisation of human DNA replication origins by initiation site sequencing (ini-seq). Nucleic Acids Res. 2016;44(21):10230–47.

24. Zhao PA, Sasaki T, Gilbert DM. High-resolution Repli-Seq defines the temporal choreography of initiation, elongation and termination of replication in mammalian cells. Genome Biology. 2020;21(1):76.

25. Mas AM, Goñi E, Ruiz de los Mozos I, Arcas A, Statello L, González J, et al. ORC1 binds to cis-transcribed RNAs for efficient activation of replication origins. Nature Communications. 2023;14(1):4447.

26. Langley AR, Gräf S, Smith JC, Krude T. Genome-wide identification and characterisation of human DNA replication origins by initiation site sequencing (ini-seq). Nucleic acids research. 2016;44(21):10230–47.

27. Boos D, Yekezare M, Diffley JF. Identification of a heteromeric complex that promotes DNA replication origin firing in human cells. Science. 2013;340(6135):981-4.

28. Kamimura Y, Tak YS, Sugino A, Araki H. Sld3, which interacts with Cdc45 (Sld4), functions for chromosomal DNA replication in Saccharomyces cerevisiae. Embo j. 2001;20(8):2097–107.

29. Tsunematsu T, Takihara Y, Ishimaru N, Pagano M, Takata T, Kudo Y. Aurora-A controls pre-replicative complex assembly and DNA replication by stabilizing geminin in mitosis. Nature Communications. 2013;4(1).

30. Cayrou C, Grégoire D, Coulombe P, Danis E, Méchali M. Genome-scale identification of active DNA replication origins. Methods. 2012;57(2):158–64.

31. Wang W, Klein KN, Proesmans K, Yang H, Marchal C, Zhu X, et al. Genome-wide mapping of human DNA replication by optical replication mapping supports a stochastic model of eukaryotic replication. Mol Cell. 2021;81(14):2975–88.e6.

32. Sansam CG, Pietrzak K, Majchrzycka B, Kerlin MA, Chen J, Rankin S, et al. A mechanism for epigenetic control of DNA replication. Genes Dev. 2018;32(3-4):224–9.

33. Tsunematsu T, Takihara Y, Ishimaru N, Pagano M, Takata T, Kudo Y. Aurora-A controls pre-replicative complex assembly and DNA replication by stabilizing geminin in mitosis. Nat Commun. 2013;4:1885.

34. Gibson DG, Young L, Chuang RY, Venter JC, Hutchison CA, 3rd, Smith HO. Enzymatic assembly of DNA molecules up to several hundred kilobases. Nat Methods. 2009;6(5):343–5.

35. Randolph LN, Bao X, Zhou C, Lian X. An all-in-one, Tet-On 3G inducible PiggyBac system for human pluripotent stem cells and derivatives. Sci Rep. 2017;7(1):1549.

36. Yusa K, Zhou L, Li MA, Bradley A, Craig NL. A hyperactive piggyBac transposase for mammalian applications. Proc Natl Acad Sci U S A. 2011;108(4):1531–6.

37. Boyd K, Bottoms C, Sansam CL. SansamLab-Pipelines-Genomics/Cut_And_Run_Analysis_SnakeMake: v1.3.2 (v1.3.2). 2024.

38. Köster J, Rahmann S. Snakemake—a scalable bioinformatics workflow engine. Bioinformatics. 2012;28(19):2520–2.

39. Bolger AM, Lohse M, Usadel B. Trimmomatic: a flexible trimmer for Illumina sequence data. Bioinformatics. 2014;30(15):2114–20.

40. Martin M. Cutadapt removes adapter sequences from high-throughput sequencing reads. EMBnetjournal. 2011;17(1):10.

41. Langmead B, Salzberg SL. Fast gapped-read alignment with Bowtie 2. Nature Methods. 2012;9(4):357–9.

42. Tarasov A, Vilella AJ, Cuppen E, Nijman IJ, Prins P. Sambamba: fast processing of NGS alignment formats. Bioinformatics. 2015;31(12):2032–4.

43. Danecek P, Bonfield JK, Liddle J, Marshall J, Ohan V, Pollard MO, et al. Twelve years of SAMtools and BCFtools. GigaScience. 2021;10(2).

44. Amemiya HM, Kundaje A, Boyle AP. The ENCODE Blacklist: Identification of Problematic Regions of the Genome. Scientific Reports. 2019;9(1).

45. Ramírez F, Ryan DP, Grüning B, Bhardwaj V, Kilpert F, Richter AS, et al. deepTools2: a next generation web server for deep-sequencing data analysis. Nucleic Acids Research. 2016;44(W1):W160–W5.

46. Zhang Y, Liu T, Meyer CA, Eeckhoute J, Johnson DS, Bernstein BE, et al. Model-based analysis of ChIP-Seq (MACS). Genome Biol. 2008;9(9):R137.

47. Lawrence M, Huber W, Pagès H, Aboyoun P, Carlson M, Gentleman R, et al. Software for Computing and Annotating Genomic Ranges. PLoS Computational Biology. 2013;9(8):e1003118.

48. Ross-Innes CS, Stark R, Teschendorff AE, Holmes KA, Ali HR, Dunning MJ, et al. Differential oestrogen receptor binding is associated with clinical outcome in breast cancer. Nature. 2012;481(7381):389–93.

49. Love MI, Huber W, Anders S. Moderated estimation of fold change and dispersion for RNA-seq data with DESeq2. Genome Biology. 2014;15(12):550.

50. Langley AR, Graf S, Smith JC, Krude T. Genome-wide identification and characterisation of human DNA replication origins by initiation site sequencing (ini-seq). Nucleic Acids Res. 2016;44(21):10230–47.

51. Mas AM, Goñi E, Ruiz De Los Mozos I, Arcas A, Statello L, González J, et al. ORC1 binds to cis-transcribed RNAs for efficient activation of replication origins. Nature Communications. 2023;14(1).

