## Supplementary figures and images for "Cell Cycle-Dependent TICRR/TRESLIN and MTBP Chromatin Binding Mechanisms and Patterns"

### Supplemental Figure 1

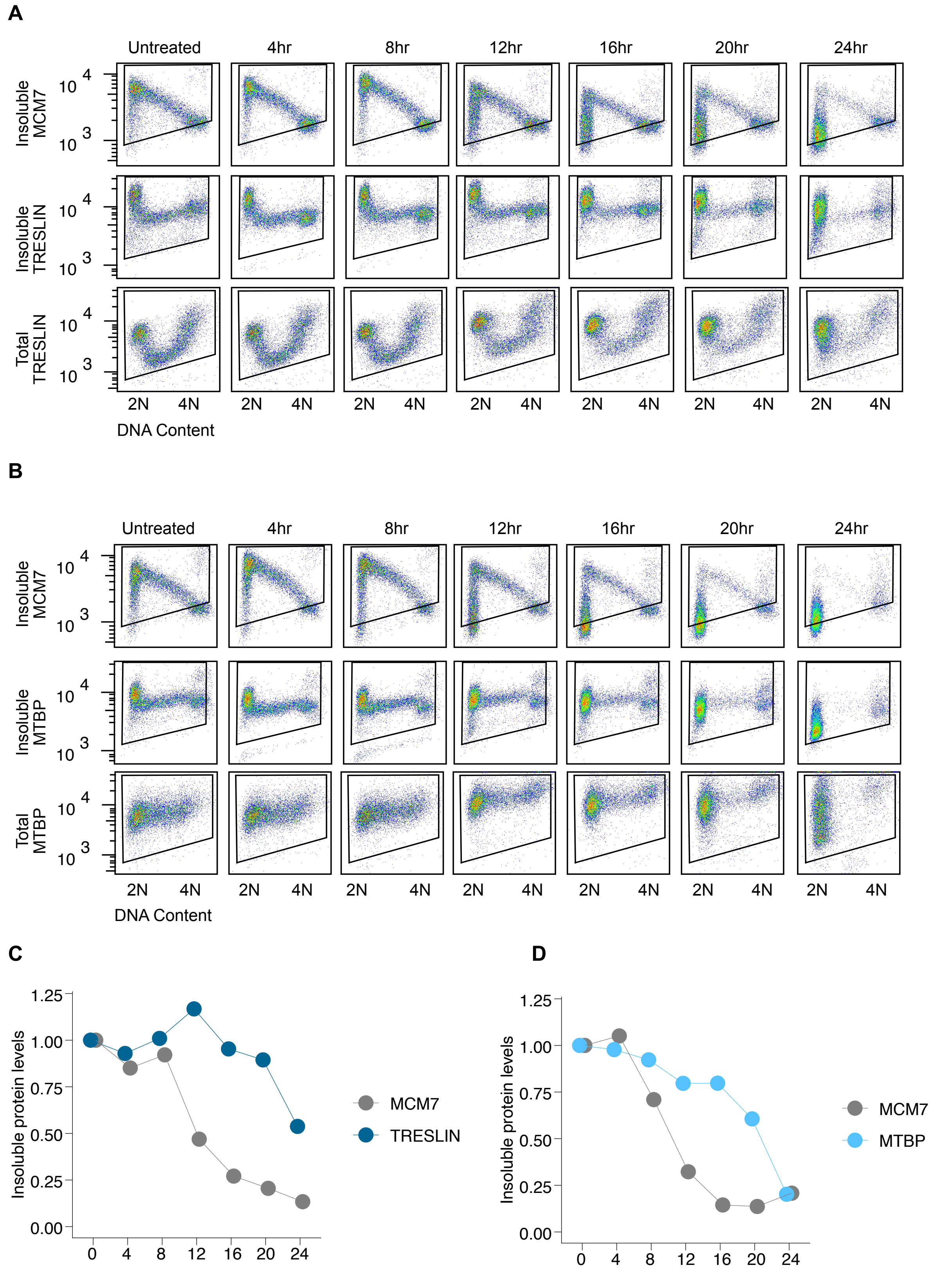
